# Macrophage phagocytosis of *Coccidioides* promotes its differentiation into the parasitic form

**DOI:** 10.1101/2025.10.05.680493

**Authors:** Jane Symington, Apoorva Dabholkar, Bevin English, Mark Voorhies, Anita Sil

## Abstract

*Coccidioides* is an endemic fungus that is increasing in prevalence and can cause life threatening disease in otherwise immunocompetent people. In the environment the spores (arthroconidia) develop into hyphae, yet when they are inhaled by a mammalian host, they develop into a unique form called the spherule. The transition to spherule can be triggered *in vitro* with elevated temperatures and high CO_2_ levels, but the signals and host cells that might trigger *Coccidioides* spherulation *in vivo* are not known. We used live imaging to investigate how macrophages affect the fate of *Coccidioides* arthroconidia. Under tissue culture conditions, arthroconidia quickly developed into hyphae. The addition of macrophages promoted spherule development and delayed hyphal formation, indicating that innate immune cells can influence *Coccidioides* development into the pathogenic form. Exposure of arthroconidia to macrophage supernatants was not sufficient to stimulate spherulation, which was dependent on phagocytosis of arthroconidia by macrophages. Transcriptomics analysis of *Coccidioides* co-cultured with macrophages revealed a signature concordant with spherules grown *in vitro* and allowed the identification of a core set of spherule-specific transcripts. In addition, we identified *Coccidioides* transcripts with significantly higher abundance in the presence of macrophages compared to *in vitro* spherules, suggesting these factors may be needed to survive and thrive in the presence of innate immune cells. This work lays a foundation for uncovering host-pathogen signaling as well as *Coccidioides* factors that are critical for pathogenesis.

## INTRODUCTION

*Coccidioides* spp. are fungal pathogens that cause Coccidioidomycosis or Valley Fever in both immunocompetent and immunocompromised individuals. Although many experience a self-limiting disease, others develop severe disseminated forms including meningitis which requires lifelong therapy (1). Coccidioidomycosis cases are rising nationally, and the endemic area may be spreading, making better diagnostics and therapeutics essential. Our understanding of this pathogen is limited in part because *Coccidioides* has a unique parasitic morphological form called the spherule. In the environment *Coccidioides* grows as hyphae, but when the spores (arthroconidia) are inhaled they undergo a morphologic transition to the spherule which can grow as large as 40-100 μm in diameter. The spherule is filled with hundreds of internal cells called endospores (2), which are released upon spherule rupture. Each endospore can develop into its own spherule and further the spread of disease.

Our understanding of spherule development is largely based on *in vitro* experiments with specialized defined media, elevated temperatures (39 °C), and elevated CO_2_ (∼10-20%) (3-5). These studies have produced key understandings of the timing of events in the development of spherules from arthroconidia (3, 6) yet the relationship of these *in vitro* studies to events within a mammalian host are unknown. Many open questions about *Coccidioides*-host interactions remain, including how the transition to spherules is triggered in the host and how arthroconidia avoid clearance by innate immune cells.

The initial infection site with *Coccidioides* is the lung. Respiratory pathogens must overcome the host defenses of the lung including macrophages, other immune cells, and epithelial cells, to cause disease. Many successful pathogens, including *Histoplasma capsulatum* and *Mycobacterium tuberculosis* subvert anti-microbial processes in macrophages to survive in the host (7, 8). Early papers suggested that a variety of immune cells could kill arthroconidia, delay germination of hyphae, or possibly promote spherules (9-11). Here we take advantage of live-cell imaging in a biosafety level 3 facility as well as transcriptomics studies to assess how macrophages affect the developmental and molecular fate of *Coccidioides* arthroconidia. We determined that murine bone-marrow derived macrophages (BMDM) stimulated a significant increase in spherule formation and a robust delay in hyphal formation in a phagocytosis-dependent manner. Using RNAseq to determine the transcriptional response of *Coccidioides* to macrophage co-culture, we observed that the fungus accumulates transcripts associated with *in vitro* spherulation, as well as a unique set of transcripts that were more highly abundant in the presence of macrophages. Together, these studies elucidate how *Coccidioides* adapts to the presence of host immune cells by forming spherules and inducing potential virulence factors to cause disease.

## RESULTS

### Bone-marrow derived macrophages *promote Coccidioides* spherule development

We used live imaging to compare the germination and subsequent development of arthroconidia in tissue culture conditions (37 °C, 5 % CO_2_, in bone marrow derived macrophage media) alone or in the presence of BMDM. In the absence of macrophages, arthroconidia primarily develop into hyphae, rarely forming spherules (Fig. 1A). Strikingly, in the presence of BMDM, the majority of arthroconidia developed into spherules (Fig. 1B, 1C). The spherules formed in the presence of macrophages were also larger in diameter than spherules formed under tissue culture conditions (Fig. 1D). In addition to the promotion of spherules, the presence of macrophages significantly delayed the time it took for hyphae to first be observed in the field, which we described as “time to first hyphae” (Fig. 1E). Since time to first hyphae was a function of both the rates of arthroconidial germination and hyphal growth, we note that either or both could have been affected by BMDM.

**Figure 1.**
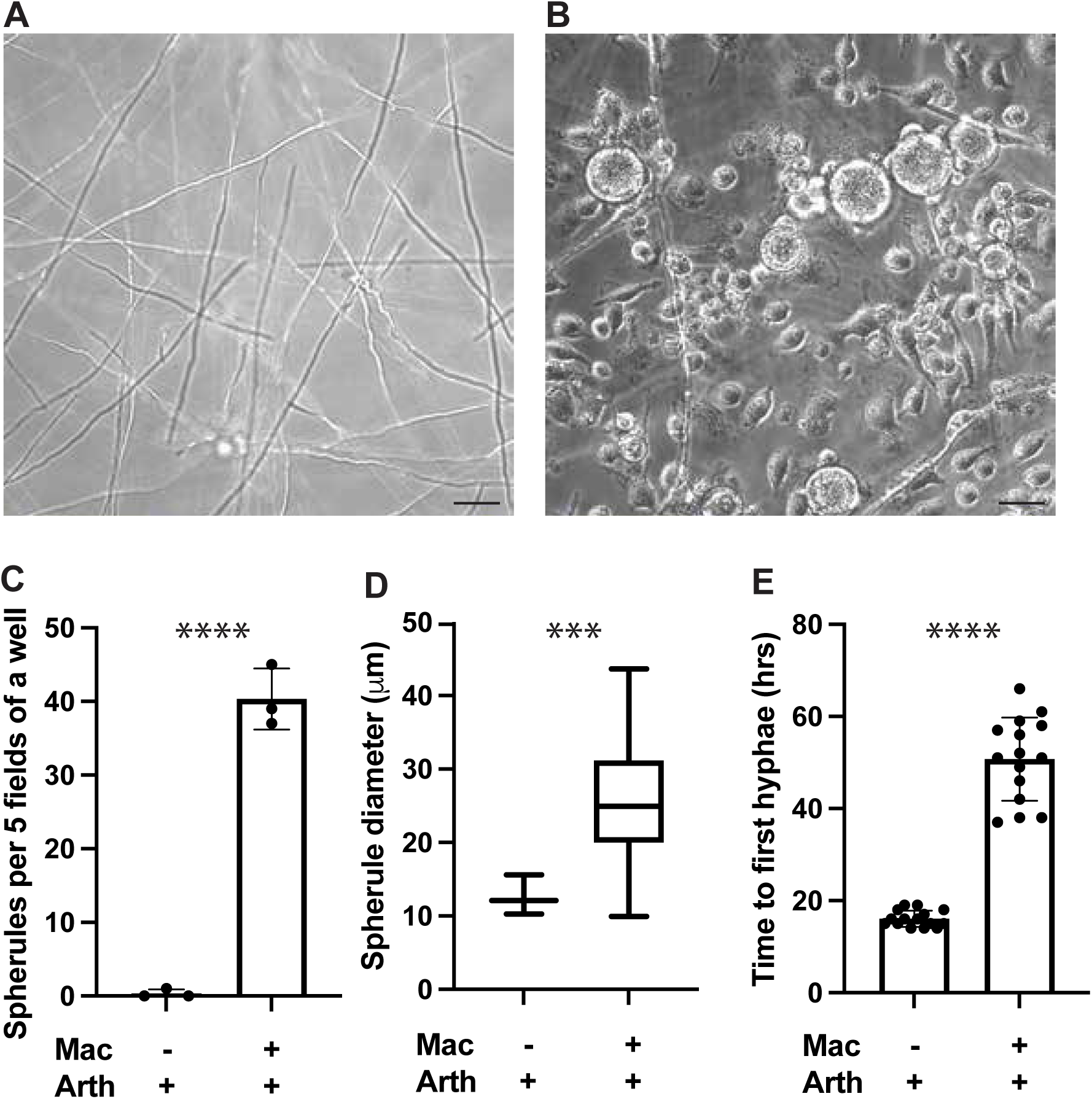
Macrophages promote spherule formation and delay hyphae. Representative image of (A) arthroconidia alone or (B) arthroconidia with BMDM at a multiplicity of infection of 0.1 (1 arthroconidia for every 10 BMDM) on day 3 of experiment, 40x (scale bar represents 30 μm). (C) Number of spherules per well (each point represents data from 5 fields per well with three independent wells per condition). (D) Average diameter of spherules on day 3 of infection in wells with arthroconidia alone or with BMDM represented as box and whisker plot. (E) Time to first hyphae in frame over 3 days with pictures taken once an hour, 5 fields per well. Data for (C, D, and E) represents 3 separate experiments, each with five 40x fields counted per well with three wells per condition. Unpaired t test used for C and Mann Whitney test for D and E, error bars in C and E indicate standard deviation, *** p < 0.001 and **** p < 0.0001.

We investigated whether these morphological findings were affected by the number of arthroconidia per macrophage (multiplicity of infection, or MOI). At the lower inoculum (MOI of 0.01), there were still significantly more spherules in the presence of macrophages than when the same arthroconidial inoculum was grown alone (Fig. 2A). Notably, the size of the spherules at MOI 0.01 was larger than the spherules that developed at an MOI 0.1 (average diameter of 54 µm vs 29 µm) (Fig. 2B). The size of the spherules that formed in the presence of macrophages was more comparable to the size of spherules observed previously *in vivo* (40-100 μm), in contrast to the smaller spherules that are characteristic of *in vitro* spherulation (10-20 μm) (2, 3, 12). At this lower inoculum, time to observation of first hyphae in the microscopy field was longer given sparser arthroconidia/rarer events, yet there was still a significant increase in time to first hyphae in the wells with macrophages compared to those without macrophages (Fig. 2C).

**Figure 2.**
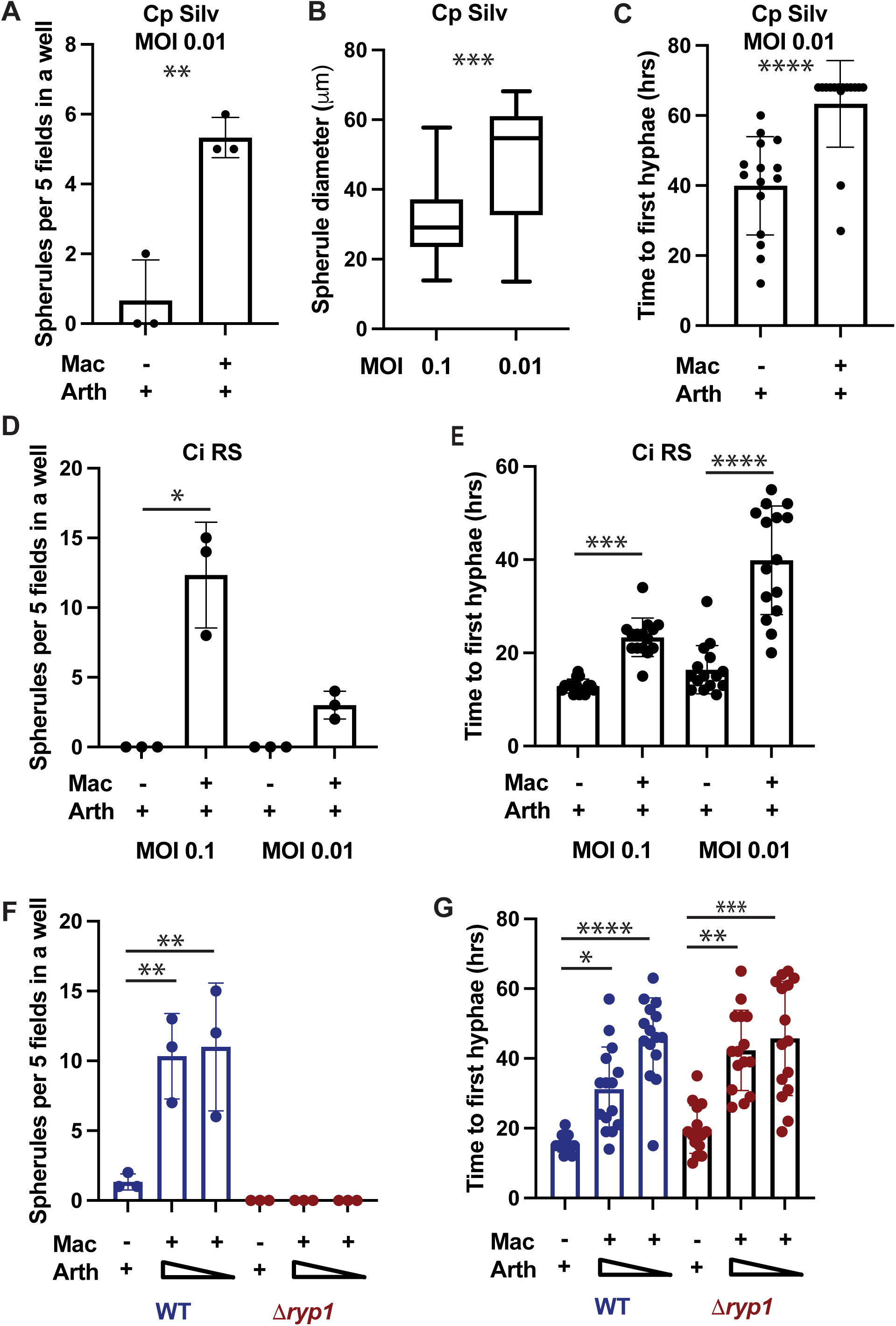
Macrophage induction of spherulation and delay of hyphae is independent of MOI and *Coccidioides* species. (A) Number of spherules per well (each point represents data from 5 fields per well with three independent wells per condition) in wells with arthroconidia alone or BMDM at MOI 0.01. (B) Average diameter of spherules on day 3 of infection at MOI 0.1 or MOI 0.01 represented as box and whisker plot. (C) Time to first hyphae in frame of arthroconidia grown alone or with BMDM at MOI 0.01. (D) Number of spherules per well and (E) time to first hyphae for *C. immitis RS* arthroconidia alone or with BMDM at MOI of 0.1 or 0.01. (F) Number of spherules per well and (G) time to first hyphae of WT and *ryp1Δ C. posadasii Silveira* arthroconidia alone or with BMDM at MOI 0.1 or 0.05. Pictures taken once an hour, 5 fields per well. Data for each figure represents five 40x fields counted per well with three wells per condition. Unpaired t test used for A, Mann Whitney test for B and C, One way ANOVA for D and F, Kruskal-Wallis test was used for E and F, * p < 0.05, ** p < 0.01, *** p < 0.001, and *** p < 0.0001.

These experiments were performed with the *C. posadasii* Silveira strain. To determine if a different *Coccidioides* species behaves similarly in the presence of macrophages, we performed the same experiments with *C. immitis*, which has more than 90% homology in predicted proteins with *C. posadasii* and a similar clinical manifestation (13). We found that macrophages also induced spherulation of *C. immitis* RS arthroconidia (Fig. 2D) and delayed the time to first hyphae (Fig. 2E).

As another contrast to the fate of wild-type *C. posadasii* Silveira, we also co-cultured macrophages with mutant arthroconidia lacking the transcription factor *RYP1*. We have previously shown that *RYP1* is essential for development of the spherule *in vitro* and in a murine infection model (6, 14). We found that *RYP1* is also essential for spherulation in the presence of macrophages (Fig. 2F). Interestingly, we found that the time to first *ryp1Δ* hyphae in the field was delayed in the presence of macrophages (Fig. 2G), implying that macrophages inhibited germination and/or hyphal growth rates of the *ryp1Δ* mutant. Since the *ryp1Δ* mutant does not make spherules, we concluded that the observed delay in time to first hyphae was not contingent on the formation of spherules by internalized arthroconidia.

### Spherule development is dependent on contact with live macrophages

Secreted compounds from macrophages could be responsible for promoting spherulation of *Coccidioides* during co-culture. To test this hypothesis, we utilized a transwell system to image arthroconidia in the lower well that either had no exposure to macrophages; had macrophages present in the same lower well, or had macrophages seeded on the insert. In the latter case, macrophages were separated from *Coccidioides* by a 0.2 μm membrane that allowed media and secreted compounds to move between wells but prevented direct contact between the arthroconidia and macrophages.

Presence of macrophages in the same wells as arthroconidia resulted in increased number of spherules per field and a delay in time to first hyphae, as expected. However, when macrophages were separated from arthroconidia by the insert, they were unable to stimulate spherulation (Fig. 3A). Additionally, the time to first hyphae in these wells was indistinguishable from wells with no macrophages (Fig. 3B). These data indicate that contact between arthroconidia and macrophages was essential for spherule stimulation and delay of hyphal appearance to be observed.

**Figure 3.**
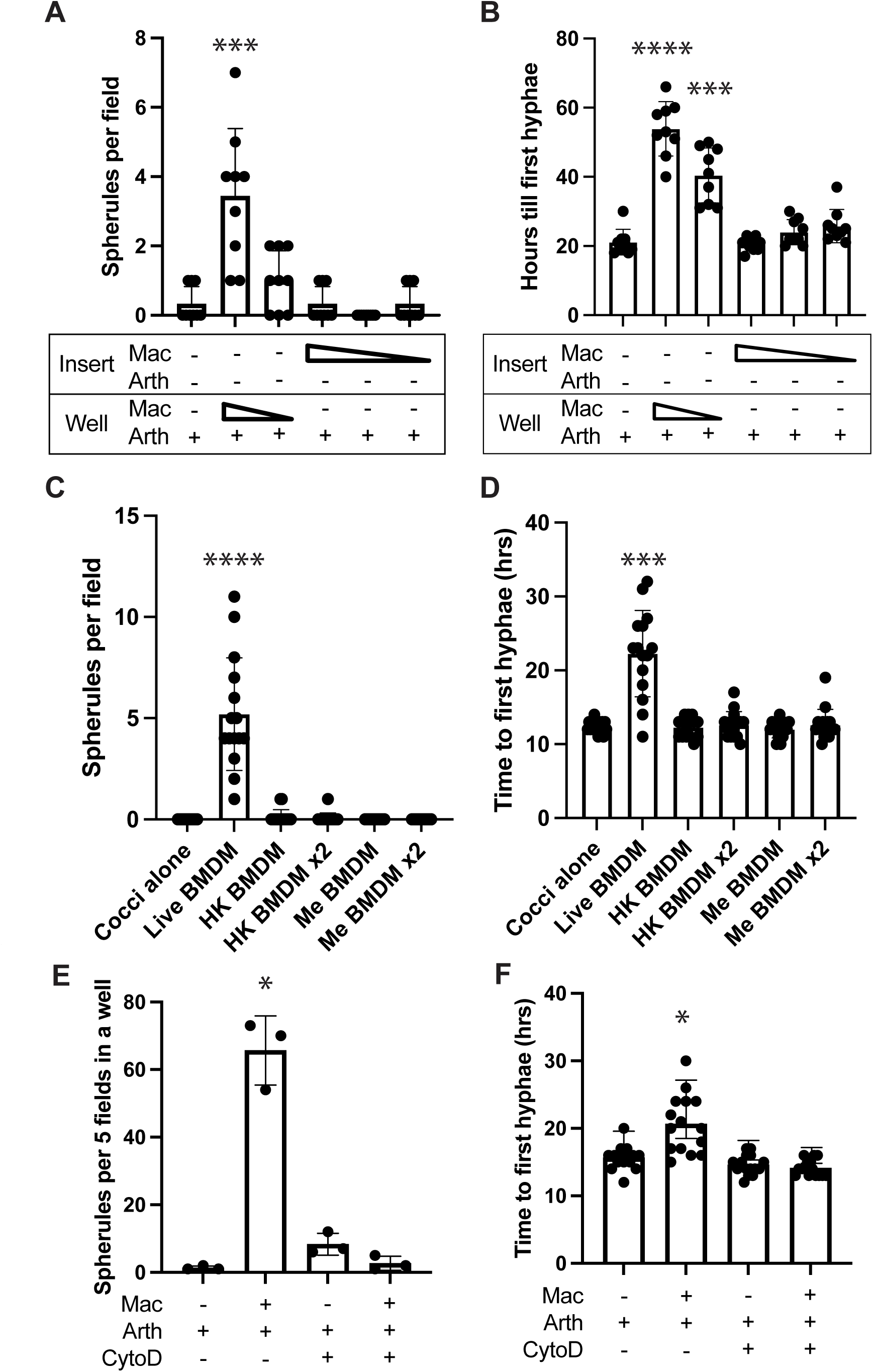
Phagocytosis of arthroconidia is required for macrophages to promote spherule formation and delay hyphae. Number of spherules per field (A) and time to first hyphae per field (B) for arthroconidia cultured with or without macrophages in the well with arthroconidia or with macrophages separated by a transwell. Number of spherules per well (C) and time to first hyphae per field (D) arthroconidia with live or heat-killed or methanol treated macrophages. Number of spherules per well (E) and time to first hyphae per field (F) of arthroconidia cultured with or without macrophages in the presence or absence of 10 μm cytochalasin D. For all samples, images were counted on day 3. For A and B, 9 fields per condition were quantified, and for C-F, 5 fields per well and 3 wells per condition were quantified. Each figure represents 3 independent experiments. Kruskal-Wallis test was used for A, B, C, D, and F, and one way ANOVA was used for E. * p < 0.05, ** p < 0.01, *** p < 0.001, and *** p < 0.0001.

Since contact with macrophages was necessary for the promotion of spherulation in our conditions, we investigated if contact with dead macrophages was sufficient to induce spherulation and delay time to first hyphae. We exposed arthroconidia to live macrophages, heat-killed macrophages, or methanol-fixed macrophages. When arthroconidia were plated with BMDM that were heat-killed or fixed with methanol, there were no changes in spherule number (Fig. 2C) or the time to first hyphae (Fig. 2D). Thus, contact with the surface of dead macrophages was not sufficient to induce spherule development.

### Phagocytosis is essential for macrophage promotion of spherulation

To test if phagocytosis was required for macrophage promotion of spherulation and delay of hyphae formation, we exposed arthroconidia to BMDM with or without cytochalasin D, an actin polymerization inhibitor that blocks phagocytosis. In the presence of untreated macrophages, arthroconidia developed into spherules at a higher rate than arthroconidia alone, but if macrophages were treated with cytochalasin D, there was no increase in spherulation compared to arthroconidia alone, with or without cytochalasin D (Fig. 2E). Similarly, the delay in hyphal formation noted in the presence of macrophages was eliminated if those macrophages were treated with cytochalasin D (Fig. 2F), suggesting that the delay in time to first hyphae was also dependent on phagocytosis.

### Co-culture with *Coccidioides* arthroconidia leads to macrophage death

Many pathogens manipulate host cell death to survive within or to defend from innate immune cells. We examined the effect of arthroconidia infection on macrophage survival by measuring release of lactase dehydrogenase (LDH) from host cells as a proxy for host-cell lysis. When macrophages were infected at an MOI of 0.1, there was no significant LDH release until day 2 of infection. When macrophages were infected at a higher MOI of 1, they showed significant LDH release at days 1, 2 and 3 post infection (Fig. S1). We concluded that co-culture with *C. posadasii* arthroconidia led to lysis of BMDM.

### *Coccidioides* induces a unique transcriptional program when challenged with macrophages

To characterize the *Coccidioides* transition to the spherule form in the presence of macrophages, we performed RNA sequencing of *Coccidioides* co-cultured with macrophages. Obtaining sufficient RNA from pathogens in co-culture can be challenging, which we overcame by utilizing multiple MOIs to increase the range of fungal burden at early timepoints, subjecting samples to bead beating to optimize lysis of fungal cells, and increasing the sequencing depth.

We examined the transcriptome of WT *Coccidioides* grown alone at 48 h, or co-cultured with BMDM at a higher MOI of 1 at 24 h (to improve RNA yield for the early timepoint) and at a low MOI of 0.1 at 24 and 48 h (to allow analysis of later *Coccidioides* with majority of macrophages still viable) (Fig 4A), each in triplicate. As above, *Coccidioides* grown alone developed into hyphae, where *Coccidioides* co-cultured with BMDM developed into spherules. To distinguish which transcriptome changes may be specific to the formation of spherules vs the delay of hyphae formation, we leveraged the *ryp1Δ* mutant, which could not make spherules in the presence of macrophages (Fig. 2F), though it did have a delay in time to first hyphae (Fig. 2G). We reasoned that transcriptome changes that correlated with spherulation might be dependent on Ryp1. Therefore, we profiled the transcriptome of the *ryp1Δ* mutant in the presence of macrophages (Fig. 4A).

**Figure 4.**
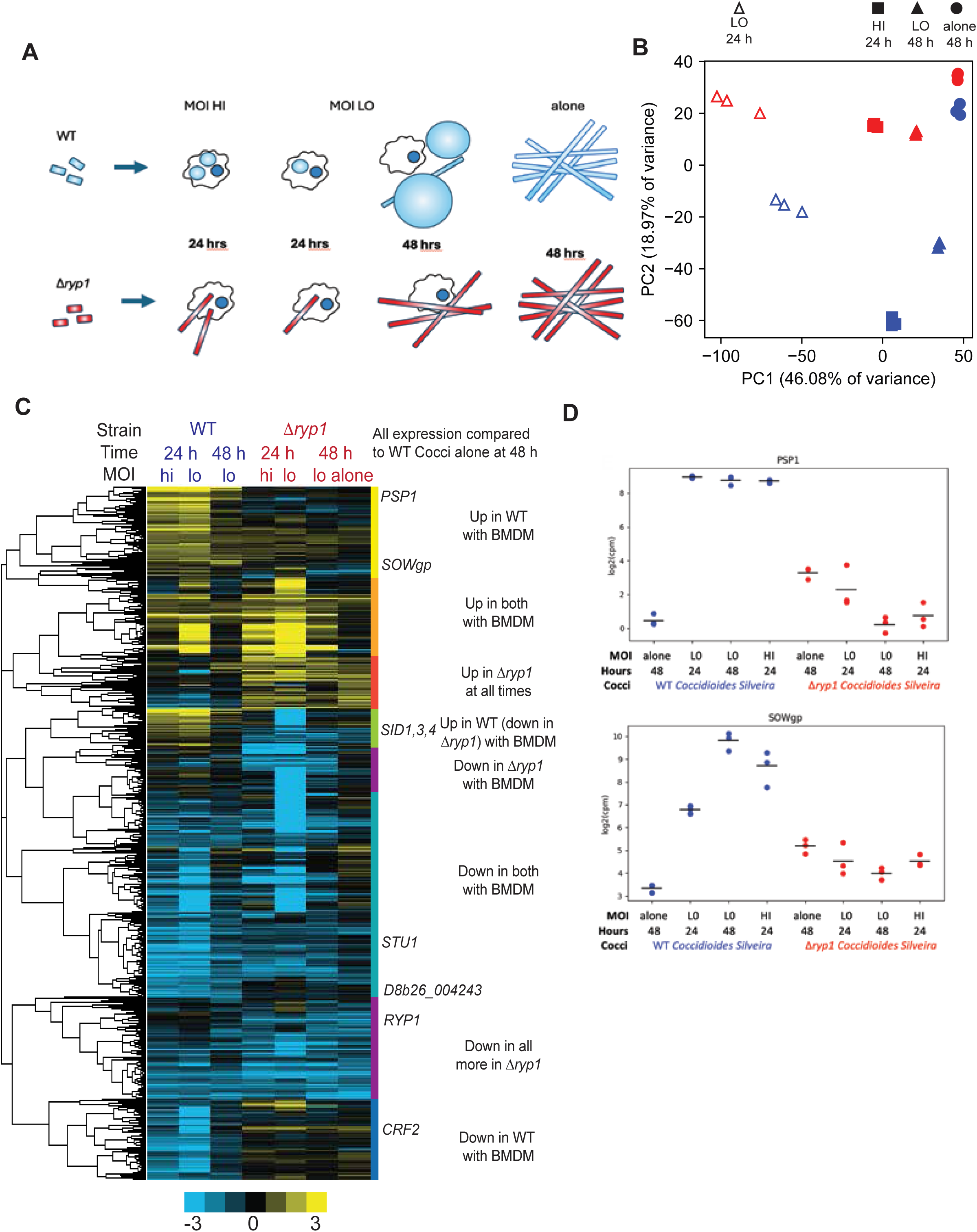
*Coccidioides* induces spherule transcriptome changes in response to BMDM. (A) Diagram of RNAseq conditions with uninfected macrophages, macrophages exposed to *ryp1Δ* arthroconidia (in red) that form hyphae but cannot form spherules or to WT *Coccidioides* (in blue) that can form spherules. “Hi” indicates MOI = 1 and “Lo” indicates MOI = 0.1. (B) PCA plot of each sample with WT in blue, *ryp1Δ* in red, Coccidioides alone at 48h is represented by circles, *Coccidioides* exposed to BMDM at lo MOI by triangles (open at 24 h and filled at 48h) and hi MOI at 24 h by squares. (C) Hierarchically clustered heatmap of differentially expressed transcripts for seven conditions, each compared to WT *Coccidioides* grown alone in tissue culture conditions. The seven conditions are WT or *ryp1Δ Coccidioides* grown in the presence of BMDM (at the indicated MOI and timepoint) or *ryp1Δ Coccidioides* grown alone under tissue culture conditions. Heatmap colors indicate log_2_ differential expression ratios relative to WT alone in units of counts per million. (D) Depth normalized abundance of transcripts (as log_2_ counts per million) of two *Coccidioides* genes known to be expressed in spherules (*PSP1* and *SOWgp*). Dots represent individual observations and bars are averages in log space.

Transcript abundance was quantified with KALLISTO and further analysis was restricted to 8274 well-represented transcripts that had at least 10 counts in at least 9 samples (Table S1). Principal components analysis of the expression profiles revealed two independent components accounting for 65% of the variance: a first component accounting for 46% of the variance that separated the samples by exposure to BMDM and time regardless of genotype (WT or *ryp1Δ*) and a second component accounting for 19% of the variance that distinguished samples with spherules from those with majority hyphae (WT *Coccidioides* with BMDM vs WT alone or vs any of the *ryp1Δ* samples) (Fig. 4B).

Differential expression analysis was carried out with the negative binomial model of edgeR to account for counting noise in the low-depth data (due to limiting fungal material in co-culture) combined with log normally distributed biological and replicate variance. For the 8274 well-sampled transcripts, we estimated contrasts for each WT and *ryp1Δ* infection timepoint and for *ryp1Δ* alone, each relative to WT alone. 3753 transcripts were at least 2-fold differential in at least one of these 7 contrasts at a false discovery rate (FDR) of 5% (Fig. 4C). Hierarchical clustering revealed 8 distinct expression patterns as indicated by the vertical rainbow color bar (Fig. 4C). More transcripts decreased in abundance in response to macrophages in both WT and *ryp1Δ Coccidioides*, compared to arthroconidia grown in the same conditions in the absence of host cells. Interestingly, prior reports of young spherules vs mature hyphal cultures grown *in vitro* also showed a smaller fraction of increased-abundance transcripts vs decreased-abundance transcripts in the young spherule (48 h) compared to mature hyphae (15).

Approximately 1/3 of differentially expressed transcripts showed increased abundance in WT *Coccidioides* in response to BMDM. These corresponded to genes previously identified as encoding spherule-associated factors, such as *SOWgp*, *PSP1*, *MDR1*, and *OPS1* (Fig. 4D)(16, 17). Other transcripts of interest that increased in abundance in WT *Coccidioides* in the presence of BMDM included *PDC1*, *UAZ1* (putative urate oxidase), and *CCC1* (a putative calcium transporter). BMDM-induced genes in WT *Coccidioides* included genes encoding hypothetical proteins of interest that were previously shown to be more significantly induced in animal models vs *in vitro* including D8B26_007421 (CPSG_01366/CIMG_09001, a putative secreted protein), D8B26_000085 (CPSG_05795/CIMG_05576, an immunoreactive protein that has been found in immunized sera from dogs that have received a live attenuated *Coccidioides* vaccine or infected with *Coccidioides*) (18), and D8B26_005342 (CIMG_00509, a putative secreted protein) (19). Thus our data suggested that the presence of macrophages could sufficiently mimic the host environment to elicit the transcription of these host-associated genes in *Coccidioides*.

Transcripts that were increased in WT *Coccidioides* exposed to BMDM but lower in *ryp1Δ Coccidioides* exposed to BMDM included D8B26_001957 (encoding a protein containing a Major Facilitator Superfamily (MFS) domain), D8B26_005576 (ortholog of iron transport multicopper oxidase *FET3*), and D8B26_000555 (a urease). Members of the siderophore biosynthesis cluster (genes *SID3*, *SID4*, *NPS1*, *OXR1*, and *MFS1* (20, 21)) were also in this group but interestingly were only significantly increased in abundance in the WT samples exposed to BMDM at 24 h but not at 48 h. This temporal regulation could be due to the local environment—at 24 h many of the WT *Coccidioides* remained within or closely associated with macrophages (potentially experiencing iron deprivation) whereas by 48 h the spherules were larger and no longer internalized.

We were interested in transcripts that were up in both WT and *ryp1Δ Coccidioides* samples in the presence of macrophages and not when either strain was cultured alone, since some of these transcripts suggested a generalized *Coccidioides* response to the stress of host cells independent of spherulation. One example was D8B26_008184, which exhibits homology to the transcription factors RlmA from *Aspergillus niger* and Smp1/Rlm1 from *Saccharomyces cerevisiae* and is proposed to be induced by cell wall stress in other fungi (22). Another gene with increased transcript abundance in all samples with BMDM was D8B26_001686, which encodes a protein of the Nramp family of metal transport proteins (similar to *S. cerevisiae* Smf1-3 transporters). D8B26_002597, which was enriched in all samples with BMDM, especially at 24 hours, appears to encode a non-ribosomal peptide synthase-like protein that might generate a natural product in response to host cells.

Since BMDM promoted *Coccidioides* spherulation and inhibited hyphal growth, we expected that hyphal-associated transcripts would show decreased abundance in WT *Coccidioides* exposed to BMDM. Indeed, genes associated with hyphal growth in *Coccidioides* and other fungal pathogens were enriched in the group of transcripts with decreased abundance in WT *Coccidioides* exposed to BMDM. One such gene was D8B26_004243, which encodes a putative Zn_2_C_6_ transcription factor that is an ortholog of *Histoplasma* Nos1, which is upregulated in hyphae, as well as *Aspergillus fumigatus* RosA and *A. nidulans* NosA, a putative regulator of sexual development (23). Transcript levels of D8B26_003897, a putative glucosidase similar to the Aspergillus fumigatus hyphal-associated glycosylhydrolase Crf2, also showed decreased abundance in WT *Coccidioides* in the presence of macrophages (24). The *Coccidioides* orthologs of two transcription factors that are key players in the hyphal program in *Histoplasma*, Stu1 (D8B26_002234) and Fbc1 (D8B26_003963) (25, 26), both showed decreased transcript abundance in WT *Coccidioides* in the presence of macrophages.

We hypothesized that the presence of BMDM induced a core set of spherule-associated genes that were also induced under *in vitro* spherulation conditions as well as a set of potential virulence factors / effectors that were specific to the presence of host cells. To identify the core genes, we leveraged our laboratory’s recent work creating a transcriptomic atlas of *Coccidioides* arthroconidia, developing spherules, and developing hyphae. We compared the gene expression data from three sets (Fig. 5A): (1) WT *Coccidioides* with BMDM (spherule) vs WT *Coccidioides* alone (hyphal); (2) *ryp1Δ Coccidioides* with BMDM (hyphal) vs *ryp1Δ Coccidioides* alone (hyphal); (3) *in vitro* spherules vs *in vitro* hyphae at day 1, 2, or 3 post-germination (6). Since sets 1 and 2 contained data from different MOIs and timepoints of BMDM infection, we considered genes to be induced if their transcripts were significantly more abundant for at least two of the three conditions in that set.

**Figure 5.**
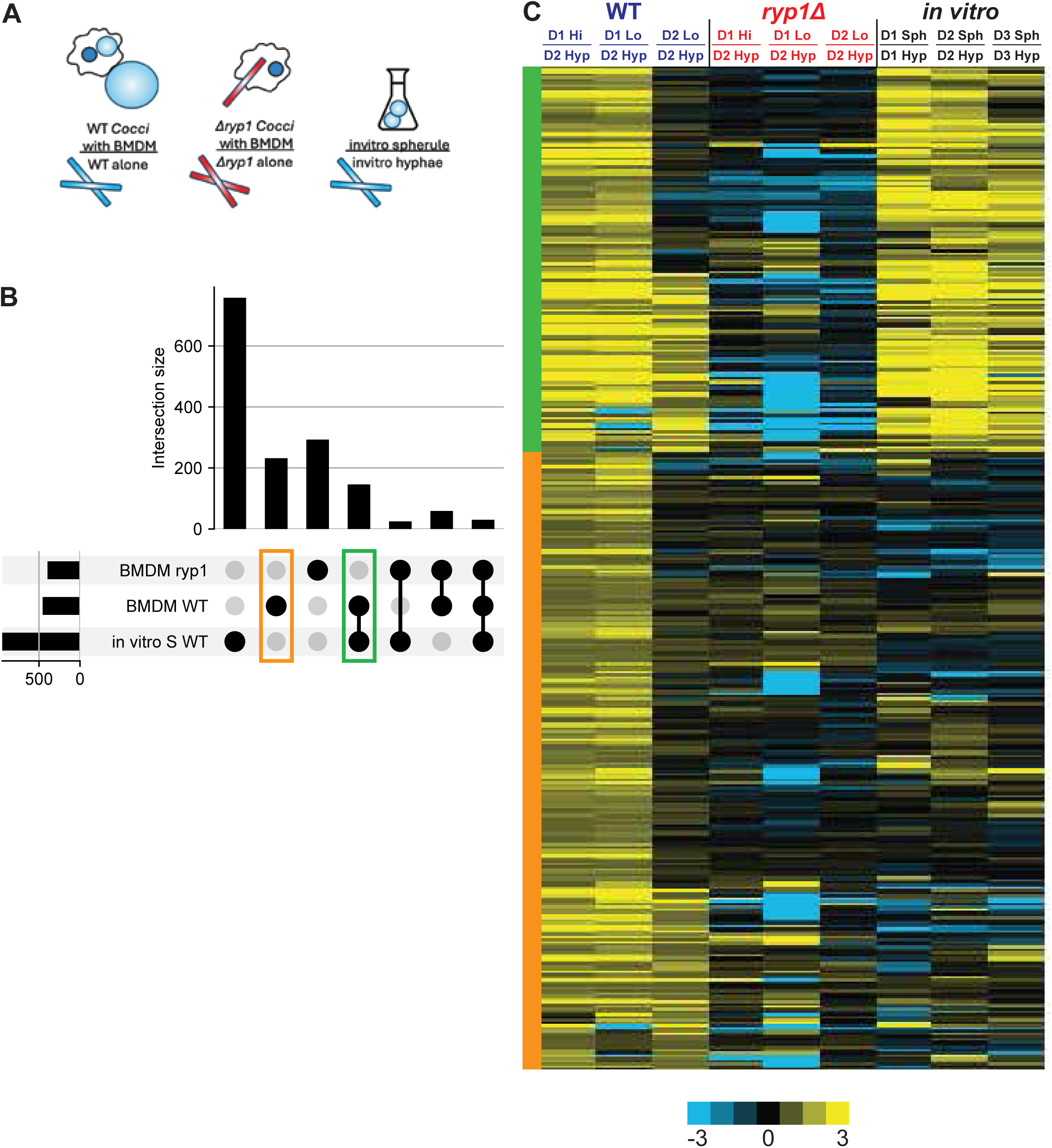
Defining core spherule associated genes and genes more significantly induced in response to BMDM. (A) Schematic of different samples used for the analysis in (B). (B) UpSet plot comparing the different gene sets depicted in (A). “BMDM WT” are transcripts with significantly increased abundance in at least 2/3 of the WT *Coccidioides* samples exposed to BMDM vs WT *Coccidioides* alone; “BMDM *ryp1*” are transcripts with significantly increased abundance in at least 2/3 of the *ryp1Δ Coccidioides* samples exposed to BMDM compared to *ryp1Δ Coccidioides* alone, and “*in vitro* S WT” are transcripts with significantly increased abundance in at least 2/3 *in vitro* spherule samples compared to *in vitro* hyphae at the same timepoint (reference Homer et al). The horizontal black bars on the left show the size of each individual set of transcripts with increased abundance as described above. The vertical black bars show the magnitude of the overlap between each of these sets (overlapping sets are indicated by connected black circles on the bottom). Each gene can only be assigned to one category per condition. Green box surrounds subset of transcripts with significantly increased abundance in both *in vitro* and BMDM-associated spherules. Orange box surrounds subset of transcripts with significantly increased abundance only in WT *Coccidioides* exposed to BMDM. (C) Heatmap of transcript abundance for WT *Coccidioides* grown in the presence of BMDM at different timepoints and MOIs vs WT *Coccidioides* grown alone at 48 h (the latter referred to as “d2 Hyp” meaning the 48 h hyphal sample); *ryp1Δ Coccidioides* grown in the presence of BMDM at different timepoints and MOIs vs *ryp1Δ Coccidioides* grown alone at 48 h (the latter referred to as “d2 Hyp” meaning the 48 h *ryp1Δ* hyphal sample); or WT *Coccidioides* grown *in vitro* as spherules at 39 °C at 10 % CO_2_ at individual timepoints compared to hyphae grown at 25 °C at the corresponding timepoints. The green and orange bars indicate the same sets of transcripts as marked in B. Heatmap colors indicate log_2_ differential expression ratios in units of counts per million.

The number of induced genes for each set and for their intersections are plotted in Fig. 5B as an UpSet plot. *In vitro* spherules had 947 differential transcripts with increased abundance in comparison to *in vitro* hyphae, the majority of which were not shared with the other contrasts, possibly due to the unique temperature and CO_2_ utilized to culture these cells *in vitro*. In the presence of BMDMs, WT and *ryp1Δ Coccidioides* had 455 and 395 differential transcripts with increased abundance compared to each of their corresponding *Coccidioides* alone controls. Although a majority of the differentially expressed transcripts in response to macrophages were unique to either WT or *ryp1Δ Coccidioides*, 83 were shared among the two. Interestingly, the shared set could represent stress-response transcripts that increased in abundance during macrophage co-culture independent of spherule formation and Ryp1 activity. More significantly, this analysis indicates that the majority of genes induced in wild-type *Coccidioides* in the presence of macrophages was dependent on Ryp1 and/or spherule formation.

We then identified the set of 143 core spherule-enriched transcripts with increased abundance in both WT *Coccidioides* in the presence of macrophages and *in vitro* spherules (green box in Fig. 5B and green bar in Fig. 5C). As proof of principle, we note this set included many of the previously identified spherule-associated genes such as *SOWgp*, *PSP1*, *OPS1*, *PDC1*, *UAZ1*, and *TYR2*. We are also highly interested in the 229 transcripts that show higher abundance only in WT *Coccidioides* in the presence of macrophages (orange box in Fig. 5B and orange bar in Fig. 5C), since these could represent virulence factors or other components required by *Coccidioides* during interaction with macrophages.

During this analysis, we noted that not all transcripts showed sustained increase in expression throughout the experiment. To further analyze expression patterns, we examined the temporal and MOI-dependent expression of the core spherule-specific genes (Fig. 5C, upper/green) and macrophage-induced genes (Fig. 5C, lower/orange). The final subsets were independently hierarchically clustered and concatenated into a single heatmap (Fig. 5C). Half of the shared enriched genes between *in vitro* spherules and WT BMDM-associated *Coccidioides* were up in all three WT-with-BMDM conditions with the other half up only in the 24 h samples. For the macrophage-induced genes, 2/3 were up only in the 24 h samples and not in the 48 h samples. This observation was interesting since, in the context of macrophage co-culture, the developing spherules tended to be intracellular at 24 h and extracellular at 48 h, suggesting that these fungal cells were in distinct environments. One of the most interesting transcripts with increased abundance in WT *Coccidioides* associated with BMDM was D8B26_000085, which encodes a highly seroreactive protein that is found in infected human lungs (19). Antibodies to D8B26_000085 persist in the serum of naturally infected or vaccinated dogs (18), indicating that it is expressed and antigenic in animal infections. Other genes with similar patterns included D8B26_005548, which contains an MFS domain, D8B26_000635, a cytochrome c peroxidase, and D8B26_006970, a presumed amidase.

To focus on potential secreted effectors that were induced when *Coccidioides* was co-cultured with macrophages, we determined which of the macrophage-induced transcripts were predicted to encode proteins with a signal peptide. 48 transcripts met this criterion, including *SOWgp*, *TYR2*, and D8B26_000085 (CPSG_05795/CIMG_05576). Next, since many fungal effectors in plant pathogens (27) and in *Histoplasma* (28) are small, cysteine-rich proteins, we further narrowed the list to small proteins that were cysteine-rich (< 250 amino acids and ≥ 4 cysteines). 10 genes encoded proteins that fit these criteria, including the superoxide dismutase *SOD3* (D8B26_006671) which is a virulence factor in *Histoplasma* (29), *ELI1* (D8B26_007114), a protective antigen of unknown function (30), two small hypothetical proteins (D8B26_007421 (CPSG_01366) and D8B26_005342 (CIMG_00509)) previously shown to be abundant *in vivo*, a predicted cutinase (D8B26_005437 which is expressed in the WT 24 h samples with BMDM), and D8B26_001315, an ortholog of *A. nidulans* calA that plays a role in conidial germination (31). These transcripts may represent key molecules at the interface between the host and pathogen during *Coccidioides* infection.

## DISCUSSION

As the endemic region for *Coccidioides* expands, putting more people at risk of disease, there is an urgent need for fundamental knowledge of how this pathogen responds to the host to cause disease. Here we showed that phagocytosis of *Coccidioides* arthroconidia by macrophages induced development of the pathogenic form of *Coccidioides*, allowing the fungus to subvert these key innate immune cells to cause disease. Furthermore, we show that *Coccidioides* exposed to BMDM expressed key genes for spherule growth as well as potential virulence factors. Lastly, we have identified a core set of spherule-associated transcripts with higher abundance in spherules, independent of temperature and CO_2_ conditions. This set will be critical for identifying targets for diagnostics or vaccine candidates in the future.

One distinguishing feature of *Coccidioides* in comparison to many other fungal pathogens is that it causes disease in immunocompetent individuals. The ability of *Coccidioides* arthroconidia to respond to macrophages by inducing spherule development and the subsequent difficulty of phagocytes to destroy the large spherule may contribute to the ability of this environmental fungus to defy the innate immune response. We have shown that after phagocytosis by macrophages, arthroconidia respond by forming spherules and that the average size of macrophage-associated spherules is larger than observed during *in vitro* spherule development. This interaction between the fungus and innate immune cells could contribute to disease progression and potentially to dissemination of the organism to extra-pulmonary sites. Notably, the specifics of the host response, which could be influenced by the initial interaction of the fungus with phagocytes, are likely to play a role in dissemination since it is known that polymorphisms in IFNγ, IL-12, and Dectin-1 are associated with the likelihood of disseminated Coccidioidomycosis (32).

This work represents the first transcriptional study of *Coccidioides* in the presence of macrophages. We overcame the technical challenges of relatively low *Coccidioides* RNA to macrophage RNA ratio with efficient disruption of fungal cells, choice of timepoints and multiplicity of infection, as well as increased depth of sequencing. One of the strengths of this study is that the conditions where WT *Coccidioides* grew as hyphae or as spherules had identical CO_2_ levels and temperature; thus differential genes in the data set reflect factors associated with spherule development and host pathogen environment rather than temperature or elevated CO_2_ alone. The identification of a core set of spherule-associated genes will be useful to our understanding of spherule biology as well the development of *Coccidioides* vaccines. We were able to observe nuance in the timing of gene expression: *Coccidioides* transcripts that are only expressed early in macrophage co-culture may reflect a germination program that gives rise to spherules, such as D8B26_001315, the ortholog of *A. nidulans* calA. D8B26_001315 is up in early WT *Coccidioides* samples exposed to BMDM as well as in early *in vitro* spherules.

Additionally, it was striking that macrophages induced the increased abundance of transcripts encoding *Coccidioides* proteins that were previously observed in human lung samples, including D8B26_000085, one of the most immunogenic proteins encoded by *Coccidioides*. These observations suggest that macrophage-*Coccidioides* co-culture provides a simplified model system for examining important host-fungal interactions.

An open question is the mechanism used by arthroconidia to sense the intracellular environment during phagocytosis to induce the spherule form. The phagosome is a complex and dynamic environment, including changes in pH, exposure to reactive oxygen species, and limitation of nutrients. What role each of those processes plays in the development of the spherule or the inhibition of hyphal germination is unknown. Furthermore, spherule formation is followed by release of internal cells named endospores, and the interaction of macrophages and endospores remains to be fully explored. An understanding of how these different *Coccidioides* cell types subvert the host immune response will provide strategies to leverage or modulate the host immune response to promote disease resolution instead of progression.

## METHODS

### Ethics statement

All mouse experiments were approved by the Institutional Animal Care and Use Committee at the University of California San Francisco (protocol AN192489-01D) and performed in compliance with the National Institutes of Health Guide for the Care and Use of Laboratory Animals). Consistent with the American Veterinary Medical Association guidelines, mice were euthanized by CO_2_ narcosis and cervical dislocation.

### Strains

Most experiments were conducted with WT *Coccidioides posadasii* strain Silveira (BEI # NR-48944), a kind gift from Bridget Barker, Northern Arizona University. We also utilized *Coccidioides immitis* strain RS (BEI # NR-48942) and *ryp1Δ* deletion mutant in the *Coccidioides posadasii* strain Silveira background (6, 14).

### Cell Culture

Bone marrow-derived macrophages (BMDM) were derived from bone marrow obtained from the femurs and tibias of 6-to-8 week old female C57BL/6 mice as previously described (33). Bone marrow cells were grown in bone marrow macrophage media (DMEM with 10 % CMG - conditioned media, 20 % FBS, Pen Strep, L glutamine, and Na-pyruvate) for 7 days then frozen for future use. For each experiment an aliquot was thawed and plated to rest overnight before each experiment. Cultures were maintained at 37 °C and 5 % C0_2_.

### Live imaging

Live imaging was performed on the Etaluma 720 microscope using the 40X objective. Cells were maintained in a stage top incubator (OKO labs) which allowed for control of temperature, humidity, and CO_2_. For most time course experiments, brightfield images of 5-9 fields per well were taken once an hour. The number of spherules per field was quantified at the final timepoint. Per well (aggregate count of number of spherules in the five fields per well) were noted. Spherule diameter was measured for all spherules in the field using Fiji. Time to first hyphae was quantified as time when the first arthroconidia germinated into a hypha in the field of view or a hypha entered the field of view.

Unless otherwise noted, experiments were performed with BMDM in 48 well plates, seeded at 0.75-1 x 10^5^ macrophages per well and infected at MOI of 0.1 arthroconidia per macrophage. Other MOIs utilized were 0.01 and 1. For transwell experiments, 6-well transwell plates were seeded with macrophages at varying densities in the well or in the transwell.

For the experiments with dead macrophages, BMDM were heat-killed at 65 °C for 5 min. For methanol fixation, BMDM were resuspended in 70 % methanol for 10 min, then spun for 5 min at 1200 rpm and resuspended in fresh BMM before plating. Cells were plated at the same density as the live BMDM (1 x 10^5^ cells per well) or at twice the density (2 x 10^5^ cells per well).

Cytochalasin D (Sigma) was used at concentrations of 10 μM in DMSO and added 30 min prior and remained throughout to infection.

### Lactate Dehydrogenase (LDH) Release Assay

To quantify macrophage lysis in response to *Coccidioides*, BMDMs were seeded (7.5 x 10^4^ cells per well) in 48-well plates and infected at MOI of 0.1 or 1 in triplicate as described above. The amount of LDH in the supernatant was measured at day 0, 1, 2, and 3 as described previously (34). BMDM lysis was calculated as the percentage of total LDH from supernatant of wells with uninfected macrophages lysed in 1% Triton X-100 at the time of infection. The total LDH at later timepoints can be greater than the total LDH from the initial timepoint due to continued replication of BMDMs over the course of the experiment. This can result in an apparent lysis that is greater than 100%.

### RNAseq Analysis

BMDM were seeded one day prior to the infection in 6-well plates. On day of infection, WT and *ryp1Δ* arthroconidia were added to the wells at an MOI of 0.1 or 1 and the plates were spun at 550 g for 5 min to bring the arthroconidia proximal to the BMDM. At 1 h, the media was removed and replaced with fresh BMM, except for the 1 h samples with BMDM and the *Coccidioides* alone wells. At each timepoint, cells were washed twice with PBS, then QIAzol was added to each of 2 wells and and incubated for 5 min at room temperature. The contents of 2 wells were then combined for each sample, with three samples per condition and per timepoint. Each sample underwent bead beating for 2 min to improve yield of fungal RNA. At this point samples were removed from the BSL3.

RNA was extracted using the Direct-zol RNA Miniprep Plus isolation kit (Zymo). Libraries were made with the NEBNext polyA mRNA magnetic isolation module and NEBNext Ultra II Directional RNA Library Prep kit with dual-indexed multiplexing barcodes as instructed. Libraries were pooled and sequencing was performed by the Chan-Zuckerberg BioHub-San Francisco on one lane of Novaseq S4.

Transcript abundances were quantified based on the annotated *C. posadasii* Silveira genome (35), PRJNA66477. Relative abundances (reported as TPM values (36)) and estimated counts (est_counts) of each transcript in each sample were estimated by alignment free comparison of k-mers between the reads and mRNA sequences using KALLISTO version 0.46.2 (37). Further analysis was restricted to transcripts with estimated counts ≥ 10 in at least nine samples.

Differentially expressed genes were identified by comparing replicates for contrasts of interest using the glmQLFit, glmQLFTest, and topTags functions in edgeR version 4.0.16 (38). Genes were considered significantly differentially expressed if they were statistically significant (at 5% FDR) with an effect size of at least 2X (absolute log2 fold change ≥ 1) for a given contrast. We estimated contrasts for each of the WT and *ryp1Δ* infection conditions, Hi MOI (1) day 1 and Lo MOI (0.1) days 1 and 2, as well as *ryp1Δ* alone all relative to WT alone. Transcripts were clustered on these 7 contrasts using maximum linkage hierarchical clustering with uncentered Pearson distances as implemented in Bio.Cluster in BIOPYTHON 1.83 (39).

To compare macrophage induced gene expression changes to those induce by *in vitro* spherulation conditions, we utilized data from Table S7 of Homer el al 2025 (6). For the *in vitro* contrasts, we contrasted arthroconidia grown in spherulation conditions (39 C, 10% CO_2_, in Converse media, shaking) and *in vitro* hyphae conditions (30 C, ambient CO_2_, in Converse media, shaking) on days 1, 2, and 3. For the with macrophages contrasts, we contrasted the gene expression of our WT and *ryp1Δ Coccidioides* in the presence of BMDMs (hi MOI 24 h, lo MOI 24 h and lo MOI 48 h) to the same strain grown alone at 48 h. Genes were considered induced in each set if they were significantly differential for at least two of the three conditions in that set.

## Figure Legends

**Figure S1.**
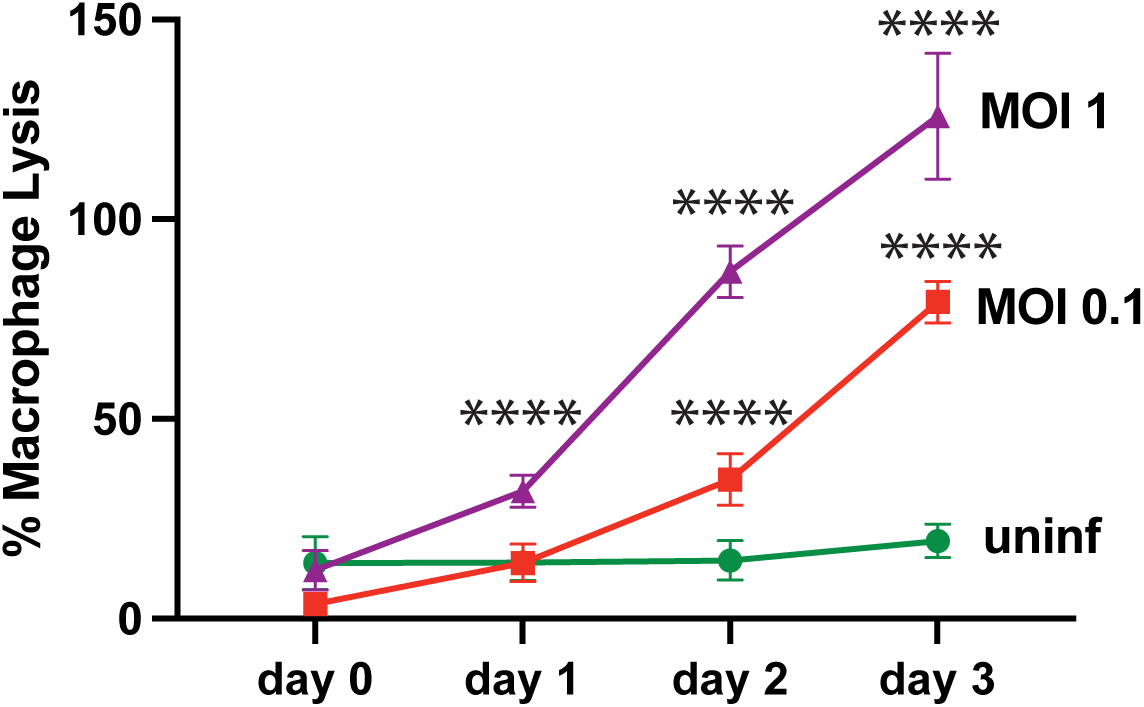
*Coccidioides* arthroconidia induce BMDM lysis. BMDM were infected with *C. posadasii* arthroconidia at an MOI of either 1 or 0.1. Lactate dehydrogenase (LDH) assay was used to measure % BMDM lysis over three days. All timepoints are relative to timepoint zero of uninfected cells lysed with 1% Triton-X (total LDH). Asterisks indicate p-value relative to uninfected according to t-test (* p ≤ 0.05, ** p ≤ 0.01, *** p ≤ 0.001.

